# Relationship between LD Score and Haseman-Elston Regression

**DOI:** 10.1101/018283

**Authors:** Brendan Bulik-Sullivan

## Abstract

Estimating SNP-heritability from summary statistics using LD Score regression provides a convenient alternative to standard variance component models, because LD Score regression is computationally very fast and does not require individual genotype data. However, the mathematical relationship between variance component methods and LD Score regression is not clear; in particular, it is not known in general how much of an increase in standard error one incurs by working with summary data instead of individual genotypes.

In this paper, I show that in samples of unrelated individuals, LD Score regression with constrained intercept is essentially the same as Haseman-Elston (HE) regression, which is currently the state-of-the-art method for estimating SNP-heritability from ascertained case/control samples. Similar results hold for SNP-genetic correlation.

## Introduction

I begin by reviewing three estimators of SNP-heritability that can be applied to GWAS data: HE regression, REML and LD Score regression. These estimators are described elsewhere, so I provide only a brief overview, with references to more detailed derivations.

Consider a model where the *N*-vector of phenotypes *Y* is generated as *y* = *X β* + *ϵ*, where *X* is an *N* × *M* matrix of standardized and centered genotypes, *β* is a vector of SNP effect sizes of length *M*, and *ϵ* is a vector of length *N* of residuals (which includes genetic effects orthogonal to an additive model, environmental effects, measurement error, etc).

If we condition on the study genotype matrix *X*, the entries of *β* as *i.i.d.* draws from a distribution with mean zero and variance 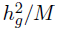, and the entries of *ϵ* as *i.i.d.* draws from a distribution with mean zero and variance 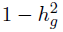, then

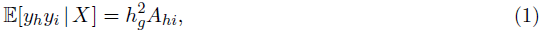

[1] where *A* is a normalized identity-by-state matrix *A* := *XX*^T^/*M.* Matrix *A* is typically called the empirical kinship matrix or genetic relatedness matrix (GRM). Equation 1 shows that we can estimate 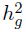 by regressing products of phenotypes *y*_*h*_*y*_*i*_ against GRM entries *A*_*hi*_ for *h < i.* This is called Haseman-Elston regression [2, 1, 3, 4]. The estimator has a closed form:

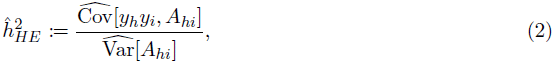

where the hats over variance and covariance denote the sample variance and covariance. The HE regression estimator is inefficient, because the datapoints are correlated; conditional on *X*, *yhyi* is correlated with *yhyj*.

If we are willing to make distributional assumptions about *β* and *ϵ*, then we can do better: if *β* and *ϵ* follow a normal distribution (or if *β* is sufficiently polygenic that *Xβ* is approximately normal by a central limit theorem argument), then *y* is distributed as 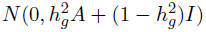, and we can estimate 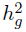 via maximum likelihood (REML) [5]. This approach is implemented in the software package **GCTA** [6](URLs).

The LD Score regression estimator of heritability [7, 8] takes as input GWAS summary statistics and LD data instead of a GRM and phenotypes. The precise LD data required are LD Scores, defined for each SNP *j* as 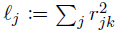, where the sum is taken over all other SNPs *k.* In practice, there is very little LD in human samples outside of small window, so LD Scores are typically estimated using a 1 centiMorgam (cM) window [7]. The GWAS summary data required are 1 degree-of-freedom *χ*^2^ statistics. Precisely, let 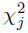 denote the Armitage Trend Test (ATT) statistic of SNP *j*, 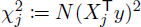 [9]. Under the same model as above, we have the regression equation

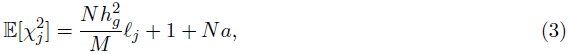

where *α* is a term that quantifies the average inflation in *χ*^2^ statistics from cryptic relatedness or population stratification [7]. We can therefore estimate heritability by regressing 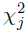 against *ℓ*_*j*_ and multiplying the slope by *M/N.* If the value of the intercept term 1 + *Na* is known ahead of time; for example, if the *χ*^2^ statistics were generated from data with relatives removed and PC covariates [10] such that *α* ≈ 0, then we can improve the efficiency of the regression can be improved by constraining the intercept. We refer to this estimator as LD Score regression with constrained intercept. The standard error can also be improved by weighting to account for heteroskedasticity [7]. Finally, the datapoints in this regression are non-independent (due to LD), so it is necessary to use a correlation-robust standard error such as a block jackknife [7, 8].

## Results

### Derivation

The HE regression estimator is typically written as a function of the GRM. However, in samples of unrelated individuals, it is possible to re-write the HE regression estimator in terms of linkage disequilibrium. Starting from the definition of covariance, we can rewrite the numerator of the HE regression estimator as

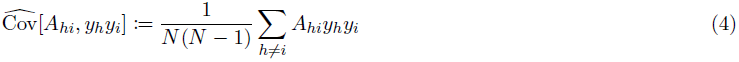

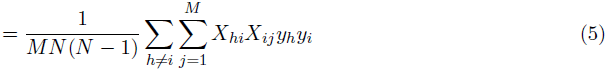

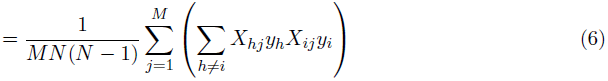

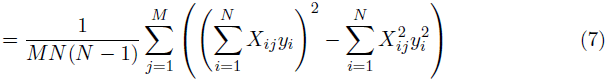

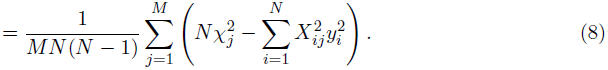

All of the preceding steps are exact and follow from the definitions of the quantities in question. I am not aware of a convenient way to simplify the term 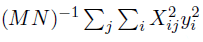 however, this term is the mean over a large number of individuals and a large number of SNPs, so the law of large numbers suggests that replacing this term with its expectation should yield a good approximation. If the marginal effect size 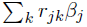 of SNP *j* is small, which is the typical case in GWAS, then *X*_*ij*_ and *y*_*i*_ will be close to uncorrelated. Even if some SNPs have large effect sizes, the average marginal variance explained will still only be 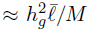, which is much less than 1 (where 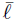 denotes mean LD Score [7]). Therefore, we approximate 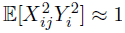. Thus,

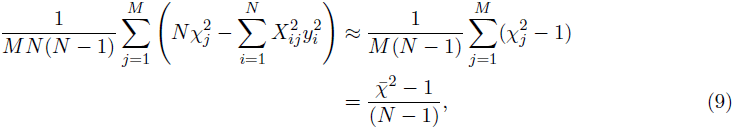

where 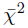 denotes mean *χ*^2^.

Next, we need to express the denominator of the HE regression estimator in terms of linkage disequilibrium. Beginning from the definition,

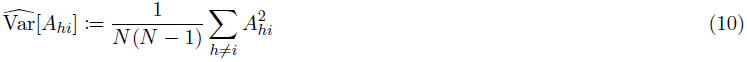

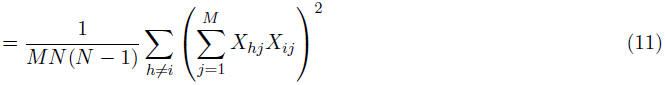

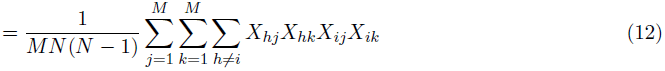

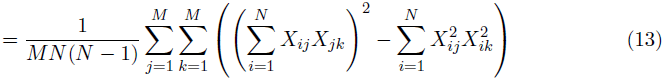

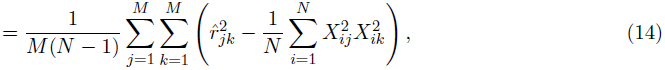

where 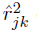 denotes the squared correlation between genotypes at SNPs *j* and *k* in our sample. The preceding steps are exact, and rely only on the definitions of the quantities in question. At this stage, we again approximate a term with its expectation. The squared sample correlation 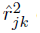 is an upwardly biased estimator the squared population correlation [11]. In fact, the bias is equal to the expectation of 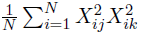. This means that the term in parentheses in Equation 14 is an unbiased estimate of the squared population correlation 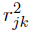. If we replace the term in parentheses with its expectation, we have

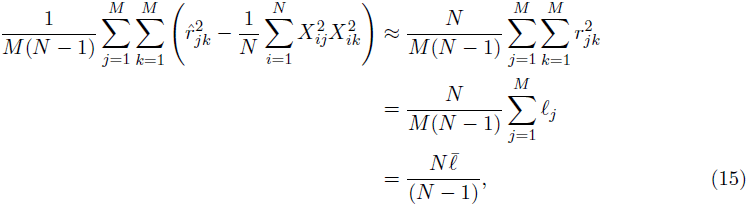

where 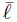 denotes mean LD Score. A similar derivation for the denominator appears in [3].

By dividing Equation 14 by Equation 15, we obtain

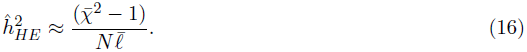

The approximation sign hides the fact that we have twice replaced terms with their expectations. However both of the terms that we replaced with expectations are means over a large number of terms, so by the law of large numbers, this approximation should be good. We verify this via simulation later in the paper.

To see that Equation 16 is equivalent to LD Score regression with the intercept constrained to one and regression weights 1/*ℓ*, first observe that by definition, unweighted LD Score regression with intercept constrained to one gives the estimator

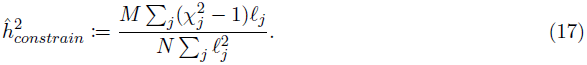

In general, weighting the regression of *y*_*i*_ on *x*_*i*_ by *w*_*i*_ is equivalent to unweighted regression of 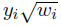 on 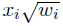. Therefore, weighting LD Score regression with constrained intercept by 1/*ℓ* is the same as regressing 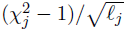 against 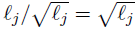 with constrained intercept. This gives the estimator

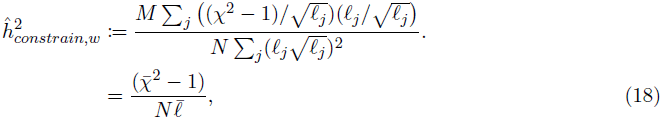

which is identical to Equation 16.

A parallel derivation in Appendix A shows that the HE regression estimator of genetic covariance is equivalent to the LD Score regression estimator of genetic covariance with 1/*ℓ*_*j*_ regression weights and in-sample LD Scores.

### Fixed Effects and Covariates

Suppose we model phenotypes as *y = Xβ* + *F* + *ϵ*, where *F* is a matrix of covariates, and *y, X, β* represent phenotypes, genotypes and effect sizes as before. Let *y′* denote *y* residualized on *F*, and let *X′* denote *X* residualized on *F.* Then the HE regression estimator of *h*^2^ controlling for fixed effects *F* is obtained by applying HE regression to *y′* and *X′.* Similarly, we can incorporate fixed effects into the ATT *χ*^2^ statistic by taking 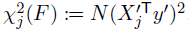. By the Frisch-Waugh-Lovell theorem [13], this *χ*^2^ statistic is equivalent to *N* times the squared standardized regression coefficient of *X*_*j*_ in the multivariate regression *y* ~ *X*_*j*_ + *F*. It then follows immediately from the previous section that HE regression with covariates is equivalent to LD Score regression with constrained intercept, 1/*ℓ* regression weights, *F*-adjusted *χ*^2^ statistics *χ*^2^(*F*), and LD Scores computed from *X′.*

What are the properties of LD Scores computed from *X′*? If *F* is a matrix of covariates that are uncorrelated with genotype in the population, *i.e.*, covariates that are not heritable, then *X′* is equal to *X* in expectation, and LD Scores computed from *X′* are equal to LD Scores computed from *X* in expectation. If *F* is a matrix of heritable covariates, then *X* and *X′* will differ in expectation. For example, if *F* is a matrix of the first 10 principal components of *X*, and *X* is a structured sample, then *X′* will be *X* with most of the population structure removed [10], and LD Scores computed from *X′* will be equal to LD Scores from *X*, except with spurious LD due to population structure removed.

The above derivations do not make any assumptions about sample structure; for example, the sample could include related individuals or a mixture of individuals with different ancestry. In these cases, HE regression is equivalent to LD Score regression with in-sample LD Scores estimated using a genome-wide window (*i.e.*, by taking the sum 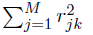 over all *M* SNPs).

The standard implementation of LD Score regression approximates in-sample LD by using LD Scores estimated from an external reference panel (such as 1000 Genomes [12]) and a 1cM window for estimating LD Scores [7]. Using a 1cM window can be viewed as a form of regularization: by taking the sum over only SNPs in a 1cM window, we reduce the variance of the estimate compared to taking the sum over all SNPs, and in samples where there is no long-range LD, we introduce only a small amount of bias. In structured samples, there will be long-range LD due to population structure; however, this LD will mostly be removed by regressing the top PCs out of the genotype matrix. If the out-of-sample LD Scores computed with a 1cM window are a good approximation to in-sample LD Scores computed with a genome-wide window after residualizing the genotypes on all principal components included as covariates in the GWAS, then the relationship between LD Score regression and HE regression should hold. If in-sample LD Score is inflated due to population structure or the inclusion of related individuals in the sample, then out-of-sample LD and in-sample LD will differ, and the estimates from HE regression and the standard implementation of LD Score regression will not be the same.

### Simulations

In order to check the approximations in the preceding derivations, I performed a series of simulations. I simulated phenotypes according to an infinitesimal model using 98,839 HapMap 3 [14] SNPs on chromosome 2 and an approximately unstructured sample of 2,062 Swedish controls from [15], chosen to be representative of a typical GWAS cohort of unrelated individuals. In all simulations, the true 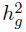 was 0.5 and effect sizes were drawn from a normal distribution. I performed 100 total simulations. I estimated heritability with five estimators: residual maximum likelihood (REML, as implemented in the software package **GCTA** [6]), HE regression, LD Score regression with unconstrained intercept and default weights (as implemented in **ldsc** [7]), LD Score regression with intercept constrained to 1 and default weights, and LD Score regression with intercept constrained to 1 and 1/*ℓ* regression weights. For all LD Score regression estimates, I used in-sample LD estimated with a 1 cM window, following [7].

Figure 1 shows a scatterplot of simulation results from all five estimators. The squared correlation between the estimates from HE regression and LD Score regression with constrained intercept and 1/*ℓ* weights in these simulations was 0.999.

**Table 1:** Comparison of Heritability Estimators. This table displays the mean and standard deviation of the 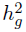 estimates from several estimators across 100 simulations of quantitative traits. The true value of 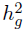 was 0.5. As expected, all estimators are approximately unbiased. REML gives the lowest standard error, followed by LD Score regression with default weights and intercept constrained to 1.

|  | Mean | SD |
| --- | --- | --- |
| REML | 0.50 | 0.050 |
| HE | 0.51 | 0.065 |
| LDSC, intercept | 0.52 | 0.091 |
| LDSC, no intercept | 0.52 | 0.060 |
| LDSC, $1/\ell$ weights | 0.52 | 0.066 |

Means and standard deviations across 100 simulations for all five estimators are displayed in Table 1. As expected, all estimators give approximately unbiased estimates. REML is the most efficient, followed by LD Score regression with default weights and constrained intercept. The worst performing estimator is LD Score regression with unconstrained intercept. Nevertheless; LD Score regression with unconstrained intercept has some advantages that may compensate for the increased standard error. Fitting an intercept protects from bias due to cryptic relatedness and population stratification [7]; however, these advantages come at the cost of an increased standard error, due to the fact that LD Score regression with unconstrained intercept fits an extra parameter.

**Figure 1:**
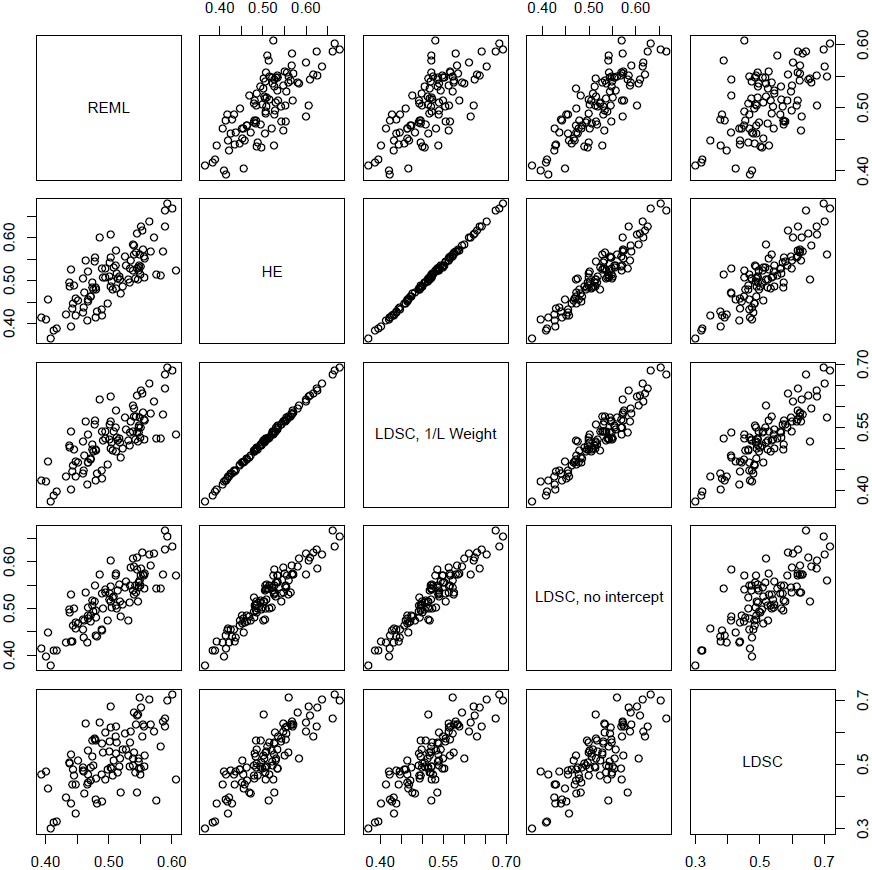
HE Regression vs LD Score Regression. Scatterplot displaying the relationships among the five heritability estimators across 100 simulation replicates. As predicted by the derivations, HE regression (HE) and LD Score regression with constrained intercept and 1/*ℓ* regression weights were almost equivalent (*R*^2^ = 0.999).

## Discussion

I have derived an approximate equivalence between HE regression and LD Score regression with constrained intercept and 1/*ℓ* regression weights. Although this equivalence is only approximate, mathematical arguments and simulations show that the approximation error is small. This provides a connection between standard kinship-based estimators of heritability and the LD Score regression estimators based on LD, and bounds the loss of precision incurred by working with summary statistics. In addition, several recent papers [3, 1] have shown that estimates of heritability from REML are biased downwards in ascertained case/control studies and recommended using HE regression instead. Since HE regression is approximately equivalent to LD Score regression, it follows that for case/control studies for which ancestry-matched LD Scores are available, LD Score regression should perform comparably to HE regression, but at lower computational cost 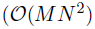 time and 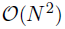 space for HE regression vs 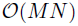 time and 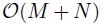 space for LD Score regression [7]).

## URLs

1. GCTA software (REML): http://www.complextraitgenomics.com/software/gcta/

2. ldsc software: github.com/bulik/ldsc

3. Coffee: http://www.trianglecoffeeshop.com

## Acknowledgements

Thanks to H. Finucane, B. Neale, M, Daly, C. Arabica, P. Sullivan, P. Fontanillas, D. Posthuma and C. de Leeuw for helpful comments.

## Appendix A Genetic Covariance

To begin, I will describe three estimators of genetic covariance that can be applied to GWAS data: HE regression, REML and LD Score regression. These estimators are derived elsewhere, so I provide only a brief overview, along with references to more detailed descriptions.

Consider a model where the vectors of phenotypes *y*_1_ and *y*_2_ are generated as *y*_1_ = *Yβ* + *δ* and *y*_2_ = *Z*_*γ*_ + *ϵ* where *Y, Z* are matrices of normalized and centered genotypes, *β, γ* are vectors of SNP effect sizes, and *δ, ϵ* are vectors of residuals (which includes genetic effects orthogonal to an additive model, environmental effects, measurement error, etc). Let *N*_1_ denote the number of individual in matrix *Y, N*_2_ the number of individuals in matrix *Z*, and *N*_*s*_ the number of individuals who appear in both matrices.

I model the entries of (*β*,*γ*) as *i.i.d.* draws from a distribution with mean zero and covariance matrix

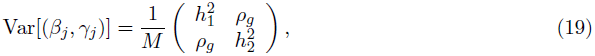

and the entries of (*δ*, *ϵ*) as *i.i.d.* draws from a distribution with mean zero and covariance matrix

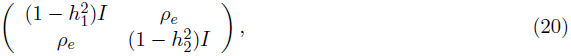

where *ρ*_*e*_ denotes the environmental covariance. who have been phenotyped for both *y*_1_ and *y*_2_. If we condition on the study genotype matrices *Y* and *Z*, the covariance matrix of the vector (*y*_1_, *y*_2_) of phenotypes is

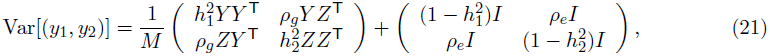

Let *A := Y*^T^*Z/M.* This means that 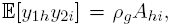, so we can estimate genetic covariance by regressing *y*_1h_*y*_2*i*_ against *A*_*hi*_. This estimator is the HE regression estimator of genetic covariance, which is inefficient for the same reasons that the HE regression estimator of heritability is inefficient.

The closed from expression for the estimator is

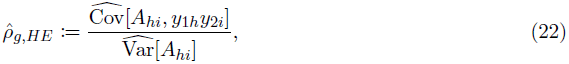

the variance and covariance are taken over all pairs (*h, i*) such that *h ≠ i*. That is, if an individual *i* is one of the *N*_*s*_ individuals phenotyped for both traits, we do not include the term *y*_1*i*_*y*_2*i*_ in the regression. However, *h ≠ i* but both *h* and *i* are among the *N*_*s*_ individuals phenotyped for both traits, we include both of the terms *y*_1*h*_*y*_2*i*_ and *y*_1*i*_*y*_2*h*_ in the regression. This is a slight difference from the single-phenotype case: if *y*_1_ = *y*_2_ then the terms *y*_1*h*_*y*_2*i*_ and *y*_1*i*_*y*_2*h*_ are identical, so it makes sense to include only one of these in the regression. If the phenotypes are not identical, then these two terms are distinct, so we would lose information by excluding one of them. There are *N*_1_*N*_2_ − *N*_*s*_ terms in the regression.

If we are willing to assume that all distributions above are multivariate normal, then the distribution of the vector of phenotypes (*y*_1_, *y*_2_) is normal with mean zero and the variance equal to the matrix from Equation 21. The REML estimator of genetic covariance is obtained by maximizing the corresponding likelihood [16].

The LD Score regression estimator of genetic covariance [8] takes as input GWAS summary statistics and LD Scores instead of a GRM and phenotypes. Let 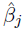 and 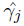 denote the estimates of the effect size of SNP *j* on *y*_1_ and *y*_2_, respectively from marginal linear regression. Under the same model from above, we have the regression equation from [8]

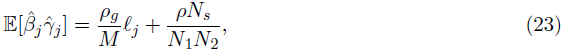

where *ρ* is the phenotypic correlation. We can therefore estimate genetic covariance by regressing 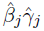 against *ℓ*_*j*_ and multiplying the slope by *M.* If the value of the intercept term *ρN*_*s*_/*N*_1_*N*_2_ is known ahead of time, the efficiency of the regression can be improved by constraining the intercept. The standard error can also be improved by weighting to account for heteroskedasticity. Finally, the datapoints in this regression are non-independent (due to LD), so it is necessary to use a correlation-robust estimator of the standard error [8].

## Genetic Covariance HE Regression and LD Score Regression

As with the heritability estimators, there is a connection between the HE regression and LD Score regression estimators of genetic covariance. If we let *S* denote the set of individuals shared by both studies, then the numerator of the HE estimator is

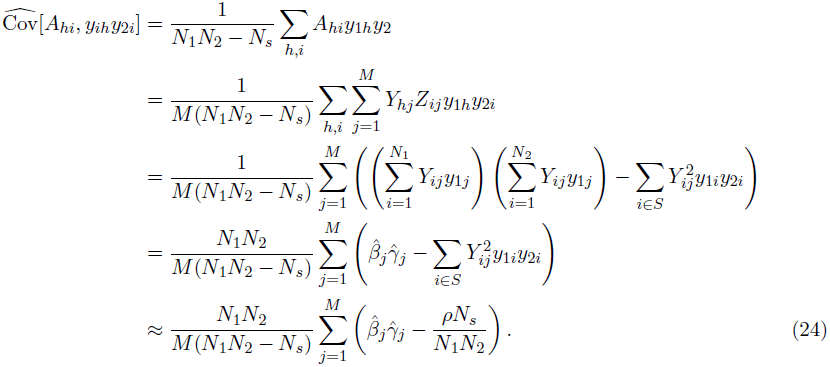

The approximation in the last line is valid in the regime of small effects, where 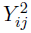 and *y*_1*i*_*y*_2*i*_ are approximately uncorrelated, in which case 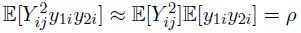.

The denominator of the HE regression estimator is

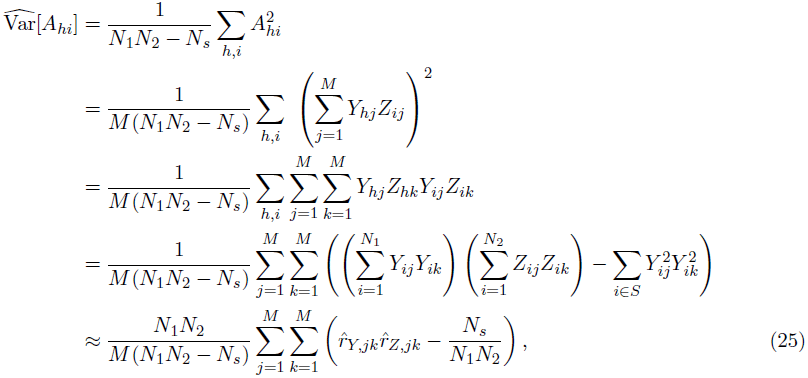

where 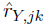 denotes the sample correlation between *j* and *k* in matrix *Y* and likewise for 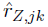. The approximation in the second-to-last line results from observing that almost all pairs of SNPs *j, k* are in linkage equilibrium, so since our genotype matrix is normalized to mean zero and variance one, then the variance of their product *Y*_*ij*_*Y*_*ik*_ will be approximately equal to the product of their variances, which is one. Observe that 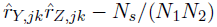 is an unbiased estimator of *r*^2^, so

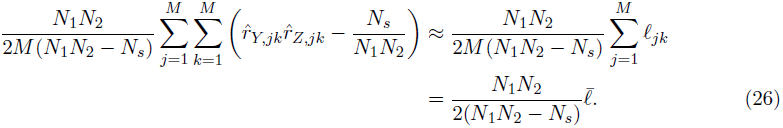

Thus, we can rewrite the HE regression estimator as

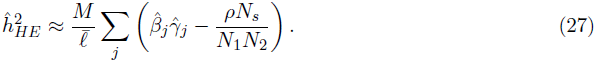

This is equivalent to LD Score regression with the intercept constrained to ρNs/(*N*_1_*N*_2_) and regression weights 1/*ℓ*. To see this, first observe that unweighted LD Score regression with intercept constrained to one yields the estimator

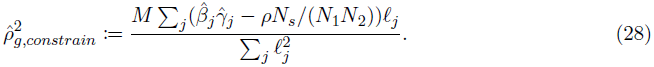

Weighting the regression by 1/*ℓ* is the same as regressing 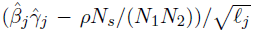 against 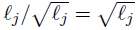 This yields the estimator

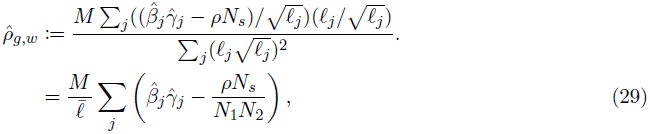

which is identical to equation 27. Using 1/*ℓ* for regression weights is more efficient than unweighted LD Score regression, but still sub-optimal [8]).

